# Chchd10 or Chchd2 are not Required for Human Motor Neuron Differentiation *In Vitro* but Modify Synaptic Transcriptomes

**DOI:** 10.1101/828376

**Authors:** Sandra Harjuhaahto, Tiina S Rasila, Svetlana M Molchanova, Rosa Woldegebriel, Jouni Kvist, Svetlana Konovalova, Markus T Sainio, Jana Pennonen, Hazem Ibrahim, Timo Otonkoski, Tomi Taira, Emil Ylikallio, Henna Tyynismaa

## Abstract

Mitochondrial intermembrane space proteins CHCHD2 and CHCHD10 have roles in diseases affecting motor neurons such as amyotrophic lateral sclerosis, spinal muscular atrophy and axonal neuropathy and in Parkinson’s disease, and form a complex of unknown function. Here we address the importance of these two proteins in human motor neurons. We show that gene edited human induced pluripotent stem cells (iPSC) lacking either CHCHD2 or CHCHD10 are viable and can be differentiated into functional motor neurons that fire spontaneous and evoked action potentials. Knockout iPSC and motor neurons sustain mitochondrial ultrastructure and show reciprocal compensatory increases in CHCHD2 or CHCHD10. Knockout motor neurons have largely overlapping transcriptome profiles compared to isogenic control line, in particular for synaptic gene expression. Our results show that absence of CHCHD2 or CHCHD10 does not disrupt functionality, but induces similar modifications in human motor neurons. Thus pathogenic mechanisms may involve loss of synaptic function.

## INTRODUCTION

Mitochondrial intermembrane space proteins CHCHD10 and CHCHD2 of unknown function have gained wide interest after being genetically associated to neurological diseases. Dominant *CHCHD10* mutations underlie motor neuron phenotypes such as amyotrophic lateral sclerosis (ALS), spinal muscular atrophy of Jokela type (SMAJ) and axonal Charcot Marie Tooth disease (CMT2) (Bannwarth *et al.*, 2014; Müller *et al.*, 2014; Auranen *et al.*, 2015; Penttilä *et al.*, 2015) whereas dominant *CHCHD2* mutations cause Parkinson’s disease (Funayama *et al.*, 2015). Evidence for whether the mutations cause disease through loss-of-function or toxic gain-of-function has been provided by different studies depending on the model system used (Genin *et al.*, 2016; Meng *et al.*, 2017; Brockmann *et al.*, 2018; Burstein *et al.*, 2018; Anderson *et al.*, 2019).

Human *CHCHD2* and *CHCHD10* originated from a gene duplication event (Cavallaro, 2010). Nuclear genes encode these small CHCH (coiled-coil-helix-coiled-coil helix) domain proteins that are imported into the mitochondrial intermembrane space by the redox-regulated MIA40 pathway (Becker *et al.*, 2012). Both proteins are conserved in metazoans (Longen *et al.*, 2009) but their exact function is unknown. They migrate in the same high molecular weight complex (Burstein *et al.*, 2018; Straub *et al.*, 2018) and have been suggested to form heterodimers (Huang *et al.*, 2018). Studies with overexpression of wild type and mutant CHCHD10 and CHCHD2 proteins, and knockdown and knockout experiments have been reported in various organisms and cultured cells, with different results depending on the mutation, cell type and the gene manipulation method. Effects on mitochondrial ultrastructure, cristae maintenance, or respiratory chain function have been identified in some (Genin *et al.*, 2016; Meng *et al.*, 2017; Huang *et al.*, 2018) but not in all studies (Burstein *et al.*, 2018; Straub *et al.*, 2018). Some studies reported that CHCHD10 is part of the mitochondrial contact site and cristae organizing system (MICOS) (Bannwarth *et al.*, 2014; Genin *et al.* 2016; Zhou *et al.*, 2018), while others did not find evidence for it (Burstein *et al.*, 2018; Straub *et al.*, 2018; Huang *et al.*, 2018). Role as a chaperone for protein import or metal transport has been proposed for CHCHD10 (Burstein *et al.*, 2018), whereas CHCHD2 has been suggested to localize also to the nucleus as a transcription factor for cytochrome c oxidase subunit 4 (COX4I2) upon oxidative and hypoxic stress (Aras *et al.*, 2015).

Knockout mice lacking CHCHD2 or CHCHD10 had a normal life span without pathological changes (Meng *et al.*, 2017; Burstein *et al.*, 2018; Anderson *et al.*, 2019). Unexpectedly, knock-in mice with the disease-associated variant p.S59L CHCHD10 developed fatal cardiomyopathy (Anderson *et al.*, 2019; Genin *et al.*, 2019). In the mice, a toxic aggregation model was proposed for pathogenesis where mutant CHCHD10 co-aggregates with CHCHD2 triggering the mitochondrial integrated stress response (Anderson *et al.*, 2019). However, a double knockout mouse model of CHCHD10 and CHCHD2 was recently introduced, which partially phenocopied the mutant S59L mouse phenotype with development of cardiomyopathy (Liu *et al.*, 2019), suggesting that the normal function of these proteins, and not just aggregates, are also relevant for pathogenesis. Nevertheless, the severe heart phenotypes in mice but not in human patients indicate that CHCHD10/2 biology somewhat differs between human and mice.

We generated knockouts of *CHCHD2* and *CHCHD10* in human induced pluripotent stem cells (iPSC) and differentiated them into motor neurons. Our results indicate that CHCHD10 and CHCHD2 are not essential for iPSC survival or for mitochondrial ultrastructure in iPSC. Although CHCHD2 was previously suggested to regulate neuroectodermal differentiation of human embryonic stem cells and iPSC by priming the differentiation potential towards neuroectodermal lineage (Zhu *et al.*, 2016), we show that human iPSC differentiate to functional motor neurons in the absence of CHCHD2 or CHCHD10. Despite being functional, we identified parallel alterations in synaptic transcriptomes of motor neurons lacking either CHCHD2 or CHCHD10, suggesting a similar function for both proteins in human motor neurons. We also observed compensatory increases in CHCHD10 or CHCHD2 that may be relevant for the development of therapeutic approaches in these motor neuron diseases.

## RESULTS

### Human iPSC without CHCHD2 or CHCHD10 are viable and pluripotent

We used CRISPR/Cas9 genome editing to generate knockouts (KO) of *CHCHD2* or *CHCHD10* in human HEL46.11 parental iPSC line (Trokovic *et al.*, 2015) by using two guide RNAs targeted to the 5’ UTR and exon 1 of these genes (Figure 1A). Four independent KO lines were obtained for *CHCHD10* (D10KO; F3, F2, B9 and G4) and one for *CHCHD2* (D2KO). Pluripotency of KO iPSC lines was verified by immunocytochemistry with Nanog and TRA1-81 antibodies (Figure 1B) and by the mRNA expression of pluripotency markers *OCT4* and *SOX2* (Figure 1C). Quantitative PCR showed complete absence of *CHCHD2* mRNA in the D2KO line and absence of *CHCHD10* mRNA in D10KO line G4, but D10KO lines F3 and F2 had 10% of residual *CHCHD10* mRNA expression (Figures 1D, S1A). Nevertheless, none of the D10KO lines had detectable CHCHD10 protein, and D2KO line had no CHCHD2 protein, as demonstrated by immunostaining (Figure 1E) and western blotting (Figure 1F). Quantification of the western blots indicated that CHCHD10 protein level was increased in the D2KO iPSC line in comparison to parental line, whereas D10KO iPSC lines had unchanged CHCHD2 protein levels (Figure 1F). We also detected higher molecular weight bands in immunoblotting with CHCHD10 and CHCHD2 antibodies that were absent in the corresponding KO cell lines (Figures S1B and S1C).

**Figure 1.**
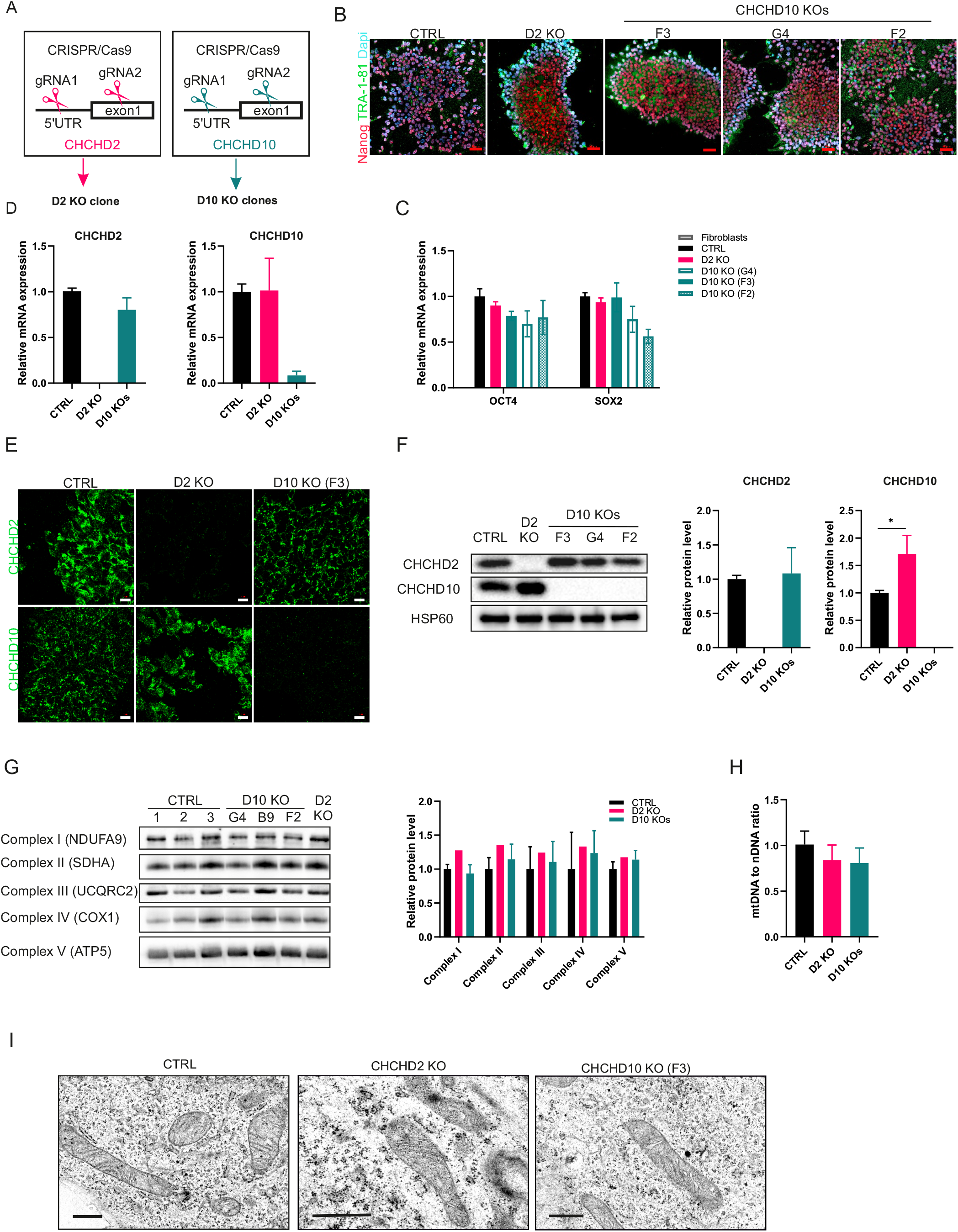
Human iPSC are viable and pluripotent in absence of CHCHD2 or CHCHD10 and have normal mitochondrial ultrastructure. (A) Schematic representation of genome editing strategy. CRISPR-Cas9 system was used to target 5’UTR and exon 1. (B) Pluripotency was validated with immunocytochemical analysis of pluripotency markers Nanog and TRA-1-81. Dapi indicates nuclear staining. Scale bar 50 μ m. (C) qRT-PCR of pluripotency markers *OCT4* and *SOX2*, normalized to *GAPDH* (n=3). (D) Expression of *CHCHD2* and *CHCHD10* in knockout iPSCs by qRT-PCR, normalized to *GAPDH* (n=3 per clone). (E) Immunocytochemical analysis verifies knockout of CHCHD2 and CHCHD10. Scale bar 20 μ m. (F) Immunoblot and quantification of CHCHD10 and CHCHD2 levels from whole cell lysates. Quantification shows the average of three independent experiments, normalized to mitochondrial HSP60. (G) Blue-native PAGE shows unaltered mitochondrial respiratory chain complexes. Quantifications shown relative to average control levels. (H) mtDNA copy number was analyzed by qRT-PCR (n=3 per clone). (I) Electron micrograph showing intact mitochondrial ultrastructure in CHCHD2 and CHCHD10 knockout iPSC. Scale bar 0.5 μ m. Quantifications of D10 KOs clones are shown as pooled results from 3 clones F3, G4 and F2. See also Figure S1 for results of individual D10KO clones and full immunoblots. Data are shown as mean ± SD, * P < 0.05.

Next, we investigated if the absence of CHCHD2 or CHCHD10 affected mitochondria in iPSC. We performed blue native polyacrylamide gel electrophoresis (BN-PAGE) to detect respiratory chain complexes and found that the overall assembly or the amount of the complexes were not consistently altered (Figure 1G). Mitochondrial DNA amount in the KO lines was comparable to the parental line (Figure 1H). Transmission electron micrographs revealed similar mitochondrial ultrastructure in KO iPSC lines as in the parental line (Figure 1I). Overall, these results suggested that CHCHD2 and CHCHD10 are not essential for iPSC survival or mitochondrial preservation in iPSC.

### Knockout iPSC differentiate into motor neurons

To investigate if human iPSC can differentiate into spinal motor neurons in the absence of CHCHD10 or CHCHD2, we followed the established differentiation protocol (Maury *et al.*, 2015; Guo *et al.*, 2017) using D2KO and three D10KO iPSC lines together with the parental control line. All lines appeared to differentiate equally well, which we verified for the neuronal differentiation by qRT-PCR of βIII-tubulin (*TUBB3*) and neurofilament medium (*NEFM*) expression (Figure 2A) and by immunostaining with NEFM, microtubule-associated protein 2 (MAP2) and βIII-tubulin (TUJ1) antibodies (Figures 2B, S3A). Differentiation towards neuronal lineage was efficient as all DAPI-positive cells were also positive for MAP2 (Figure 2B) or TUJ1 (Figure S3). Motor neuronal identity of differentiated neurons was validated by analyzing the expression levels of ISL LIM homeobox 1 (*ISL1*) and choline acetyltransferase (*CHAT*) by qRT-PCR (Figure 2A), and by immunostaining with ISL1 and motor neuron and pancreas homeobox protein 1 (HB9)/ Homeobox protein HB9 antibodies (Figure 2B). *ISL1* mRNA level was higher in D2KO neurons in comparison to parental line (Figure 2B), however, we did not observe the same in RNA sequencing of another set of differentiated neurons, showing that loss of CHCHD2 does not consistently lead to increased ISL1. Approximately 50% of DAPI-positive neurons showed pronounced HB9-positivity across all lines in each imaged frame (Figure 2C). Furthermore, we confirmed by immunostaining that motor neuron markers SMI-32 and ChAT were expressed (Figure S3B).

**Figure 2.**
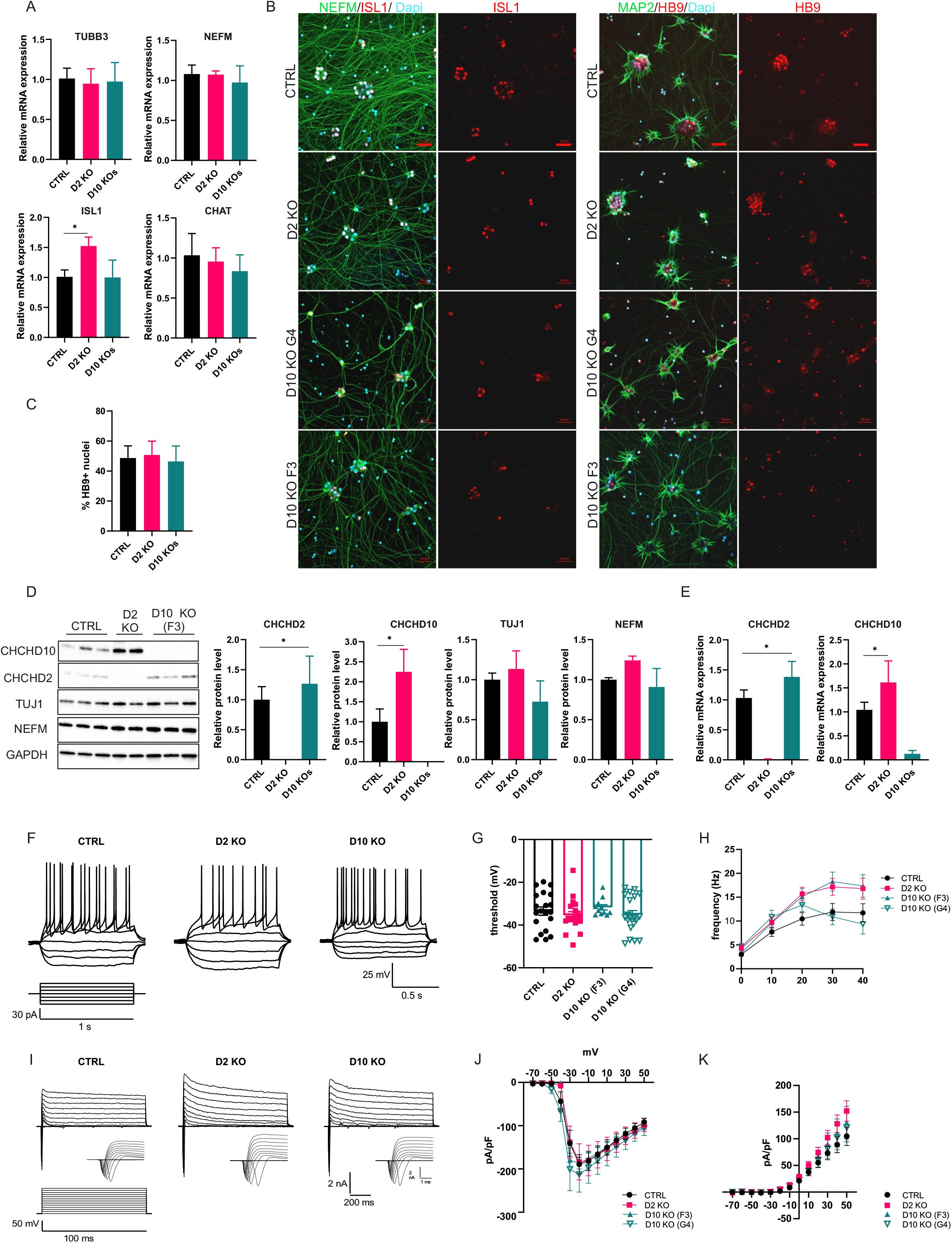
CHCHD2 and CHCHD10 knockout iPSC differentiate into functionally active motor neurons. (A) Validation of the expression of neural transcripts *TUBB3* and *NEFM* and motor neural transcripts *ISL1* and *CHAT* by qRT-PCR, normalized to *GAPDH*, on the day 37 of motor neuron differentiation (n=3 per clone). (B) Immunocytochemical analysis of ISL1 (red), NEFM (green), MAP2 (green) and HB9 (red) in neuronal cultures. Scale bars 50 μ m. Dapi indicates nuclear staining. Manual counting of HB9-positive nuclei in relation to Dapi-positive cells. Total cells counted in 3 frames (n=52-185 per clone). (D) Immunoblot and quantification of CHCHD2, CHCHD10, TUJ1 and NEFM from whole cell lysates. Quantification shows the results from two or three cultures per clone from two independent experiments, normalized to GAPDH. (E) Expression of *CHCHD2* and *CHCHD10* in knockout iPSC-derived motor neurons by qRT-PCR, normalized to *GAPDH* (n=3 per clone). (F) Representative traces of evoked action potentials of iPSC-derived motor neurons in response to current injections. (G) Threshold (mV) of action potential firing (n=11-20 per clone). (H) Action potential firing frequency (Hz) in response to incremental current injections (n=11-20 per clone). (I) Representative traces of Na^+^/Ca^2+^ and K^+^ currents, recorded at different membrane potentials. Na^+^/Ca^2+^ currents are also shown in inserts, using expanded time scale. (J) Current/voltage relationship characteristics of persistent K^+^-current normalized to membrane capacitance (pA/pF) (n=7-15 per clone). (K) Current/voltage relationship characteristics of Na^+^/Ca^2+^-current normalized to membrane capacitance (pA/pF) (n=7-15 per clone). Quantifications of D10 KOs clones are shown as pooled results from 3 clones F3, G4 and F2. See also Figure S2 for full immunoblots. Data are shown as mean ± SD, except mean ± SEM in G-K, * P < 0.05.

Next, we used immunoblotting to detect the levels of CHCHD10 and CHCHD2 in KO motor neurons. We observed a modest increase of CHCHD2 in D10KO motor neurons (Figure 2D), and a larger increase of CHCHD10 in D2KO motor neurons (Figure 2D). Higher molecular weight bands that were absent in D10KO iPSCs were again detected by immunoblotting with the CHCHD10 antibody (Figure S2A), and these were also increased in D2KO neurons. The mRNA expression level of *CHCHD10* was also increased in D2KO neurons and *CHCHD2* expression in D10KO neurons (Figure 2E). These results showed that human iPSCs differentiate successfully into motor neurons in the absence of CHCHD2 or CHCHD10, with reciprocal compensatory elevation of the two proteins. In particular, the increase of CHCHD10 was evident in CHCHD2 deficiency.

### Knockout motor neurons fire action potentials

To evaluate whether the KO motor neurons were functional and had normal electrophysiological properties, we tested them using whole-cell patch clamp during seventh week of differentiation. Current-clamp mode recordings of iPSC-derived motor neurons demonstrated that all cell lines developed the ability to fire evoked action potentials in response to current injections (Figure 2F). Many neurons also fired action potentials spontaneously, with similar frequency and amplitude regardless of cell line (Table 1). We found no differences in membrane potential threshold or frequency of firing in response to current injection (Figure 2G, H). The waveform of individual action potentials was also similar in all cell lines (Table 1). To investigate the Na^+^ and K^+^ currents in iPSC-derived motor neurons, we applied a series of depolarizing steps while recording in a whole-cell voltage clamp mode (Figure 2I). No differences in the current-voltage properties of Na^+^/Ca^2+^-currents (Figure 2K) or K^+^ persistent current (Figure 2J) were observed after normalization to membrane capacitance (pA/pF). This indicated that loss of CHCHD10 or CHCHD2 did not affect excitability of motor neurons. We also evaluated passive membrane properties in voltage clamp mode. Whole-cell capacitance (*C*_m_) values were similar across parental control and knockout iPSC-derived motor neurons, except in D10KO clone G4, which showed a reduction (Table 1). We found no significant differences in the input resistance (*R*_in_) of motor neurons. Analysis of passive membrane properties and parameters of action potential firing indicated that iPSC-derived knockout iPSC matured into functional neurons similarly to parental control. This demonstrated that functional motor neuron maturation was not compromised by the loss of CHCHD2 or CHCHD10, and thus these proteins were dispensable for human motor neuron differentiation *in vitro*.

**Table 1.** Passive membrane properties and action potential firing in motor neurons.

|  | parental control<br>(n) | D2KO (n) | D10KO F3 (n) | D10 KO G4 (n) |
| --- | --- | --- | --- | --- |
| capacitance, pF | 28.8±3.1 (25) | 22.0±1.2 (26) | 28.1±1.6 (11) | 20.5±1.8 (21)* |
| input resistance,<br>MΩ | 747.8±93.1 (25) | 827.0±90.7 (26) | 751.7±110.6 (11) | 958.2±135.8 (21) |
| resting<br>membrane<br>potential, mV | -43.2±2.0 (24) | -41.4±1.7 (22) | -48.0±1.8 (10) | -42.9±1.7 (16) |
| spontaneous AP<br>frequency, Hz | 2.2±0.4 (18) | 2.6±0.5 (13) | 1.1±0.3 (8) | 2.8±0.6 (14) |
| spontaneous AP<br>amplitude, mV | 60.0±3.8 (18) | 54.5±2.4 (13) | 63.4±3.9 (8) | 53.4±1.6 (14) |
| rheobase, pA | 24.8±3.8 (20) | 19.0±1.9 (19) | 27.3±4.5 (11) | 18.5±2.2 (20) |
| evoked AP<br>amplitude, mV | 59.7±3.2 (20) | 58.2±2.9 (19) | 56.9±2.1 (11) | 59.6±2.3 (20) |
| evoked AP half-<br>width, ms | 1.4±0.08 (20) | 1.3±0.06 (19) | 1.2±0.08 (11) | 1.2±0.04 (20) |
| after-<br>hyperpolarization<br>amplitude, mV | 17.5±2.1 (20) | 15.8±1.6 (19) | 18.1±1.1 (11) | 16.7±1.7 (20) |
| after-<br>hyperpolarization<br>decay time, ms | 13.0±0.9 (20) | 13.2±1.0 (19) | 13.5±1.9 (11) | 11.5±0.6 (20) |

n=number of cells tested. \*p<0.05 compared to parental control.

### Mitochondrial ultrastructure is preserved in knockout motor neurons

We used transmission electron microscopy to examine how the absence of CHCHD2 or CHCHD10 affected mitochondrial ultrastructure in human neuronal processes. We found abundant mitochondria in all imaged neurites in each cell line, and mitochondrial ultrastructure was similar in KO motor neurons in comparison to the parental control (Figure 3A). No significant differences in the length of mitochondria between parental control and D2KO or D10KO motor neurons were observed (Figure 3B). We categorized the mitochondria into three classes based on their length: short (<1 μm), medium (1-2 μm) and long (>2 μm) mitochondria. Most of the measured mitochondria were medium length in all studied motor neurons (Figure 3C). We determined the percentage of short, medium and long mitochondria within each cell line and found no differences in the length distribution. These results indicated that loss of CHCHD2 or CHCHD10 does not affect mitochondrial ultrastructure, abundance or length in human neurites.

**Figure 3.**
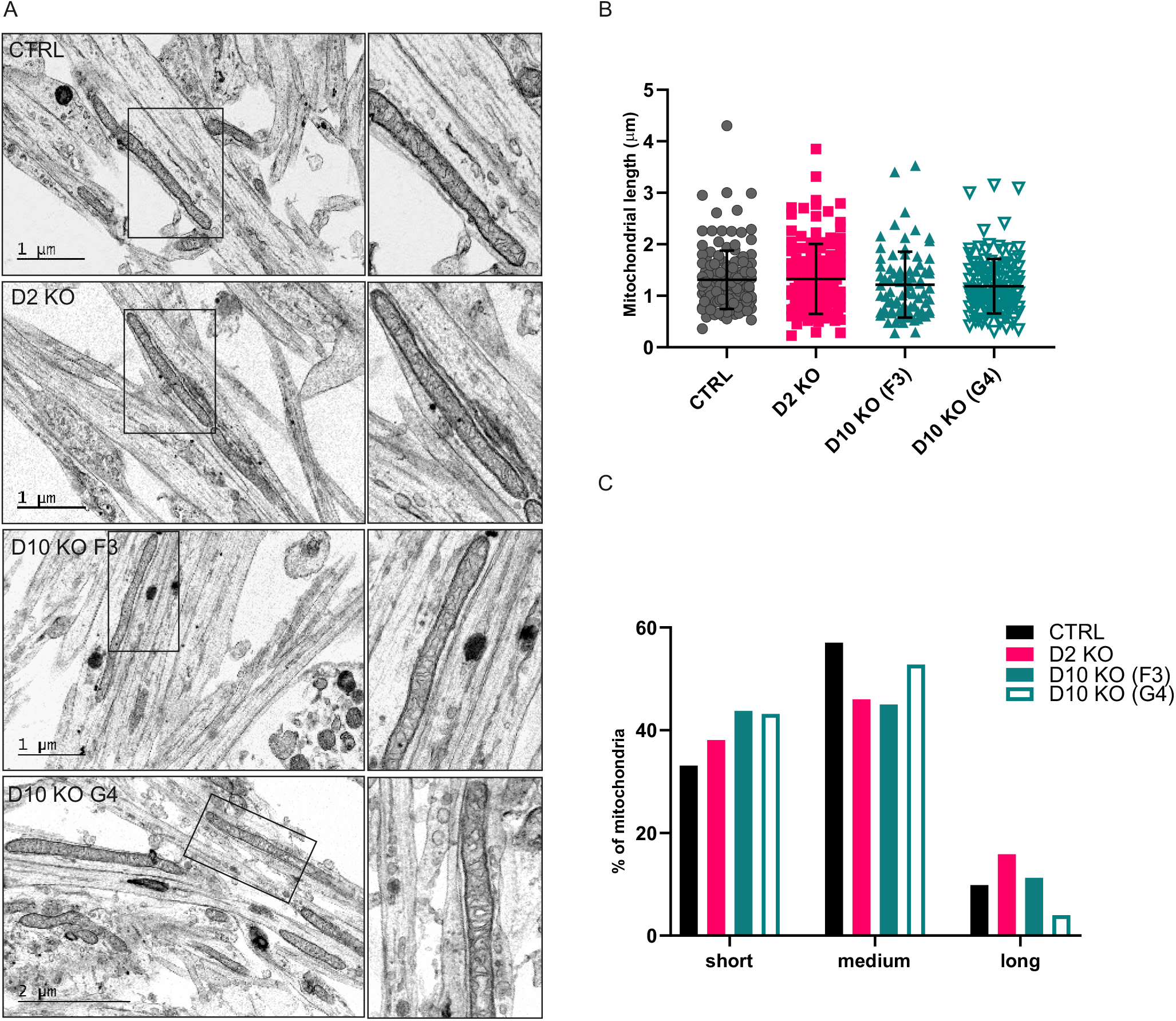
Mitochondrial ultrastructure is intact in knockout iPSC-derived motor neurons. (A) Ultrastructure analysis by transmission electronic microscopy of parental control, CHCHD2 KO and two CHCHD10 KO (clones F3 and G4) iPSC-derived motor neurons at day 38. The right panel of each image is a magnification of the area indicated by the square in the left panel. Representative images of motor neuron neurites are shown. Scale bars are indicated. (B) Quantification of mitochondrial length (μ m) in electron micrographs (n=80-143 per clone). Data are shown as mean ± SD. (C) Mitochondria length was categorized into three classes: short (<1 μ m), medium (1-2 μ m) and long (>2 μ m). Bar indicates the percentage of short, medium or long mitochondria within all measured mitochondria in each cell line.

### CHCHD10 and CHCHD2 knockout motor neurons have overlapping transcriptome profiles

Next, we assessed whether the loss of CHCHD2 or CHCHD10 had any effect on global gene expression in iPSC-derived motor neurons. We analyzed mRNA expression of four independent D2KO motor neuron cultures, four D10KO (clone G4) motor neuron cultures and eight independent cultures of the parental control motor neurons. Comparison of our parental motor neuron RNA sequencing dataset by principal component analysis (PCA) to available human iPSC-motor neuron datasets (GSE108094 (Rizzo *et al.*, 2019), GSE121069 (Nijssen *et al.*, 2018)) and laser captured post mortem human motor neuron datasets (GSE76220 (Batra *et al.*, 2016), GSE76514 (Nichterwitz *et al.*, 2016), GSE93939 (Allodi *et al.*, 2019)) showed tight clustering of our 8 samples together, supporting consistent, high-quality differentiation conditions of our cultures (Figure 4A). Interestingly, PCA of all of our samples alone showed that the expression profiles of D2KO and D10KO motor neurons complete overlapped and were separate from parental control motor neurons (Figure 4B).

Differential gene expression (DE) analysis by DESeq2 was used to compare D2KO and D10KO motor neuron transcriptomes to those of parental control. Motor neurons from both KO lines had ∼3000 differentially expressed genes (false discovery rate (FDR) < 0.01) compared to parental control motor neurons, and the majority of these changes (>65%) were shared between the knockouts (Figure 4C, Table S3). In contrast, only 175 genes were differentially expressed between D2KO and D10KO samples. These results indicated that knockout of either CHCHD2 or CHCHD10 produced similar effects on motor neuron transcriptome.

**Figure 4.**
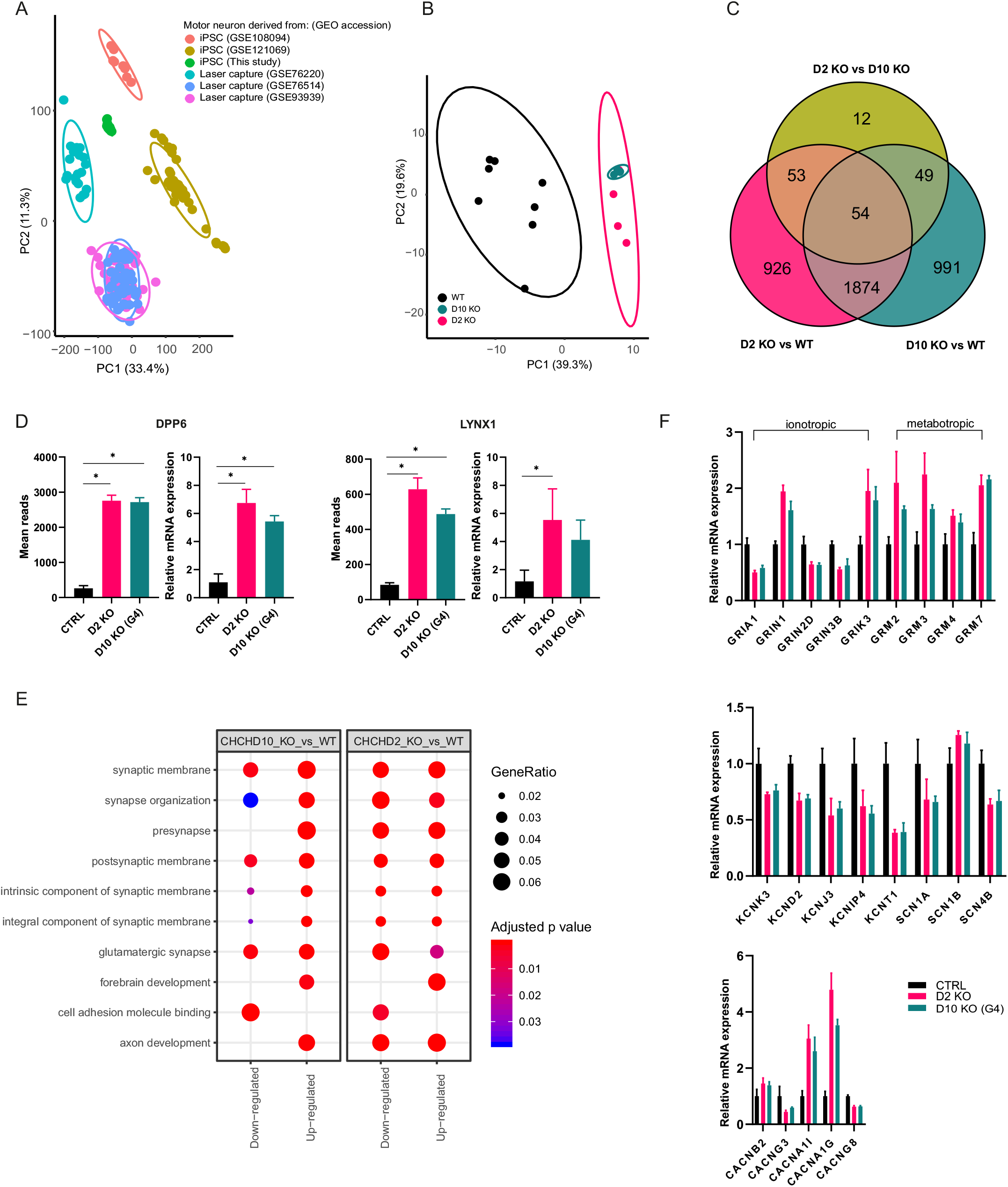
CHCHD2 and CHCHD10 knockout motor neuron transcriptome profiles are overlapping and show altered synaptic gene expression. (A) Comparison of our RNA sequencing dataset by principal component analysis (PCA) to available human iPSC-motor neuron and laser captured post mortem human motor neurons. (B) PCA of motor neurons of this study show overlap for D2KO and D10KO motor neurons (n=8 for parental control (WT) and n=4 for each KO). (C) Venn diagram displaying the number of significantly altered genes (FDR<0.01) in D10KO vs. WT, D2KO vs. D10KO and D2KO vs. WT samples. (D) Mean reads of *DPP6* and *LYNX1* in RNA sequencing dataset (left panels), and expression of the same genes by qRT-PCR (right panels) from independent samples (n=3 per clone). (E) GO term enriched pathways in D10KO vs. WT and D2KO vs. WT comparisons. Up- and downregulated pathways are indicated by gene ratio and FDR. (F) Significantly altered (FDR<0.01) glutamate receptor (upper panel), K^+^-and Na^+^-channel (middle panel) and voltage-gated Ca^2+^-channel (lower panel) subunits. Mean reads from RNA sequencing were normalized to parental control level, shown as relative expression. Data are shown as mean ± SD, * P < 0.05.

Among individual genes that were highly and significantly upregulated in both KO motor neurons in comparison to parental control motor neurons were dipeptidyl peptidase like 6 (*DPP6*) (log_2_FC = 3.5, FDR = 4.4E-103) and Ly6/neurotoxin 1 (*LYNX1*) (log_2_FC = 2.95 in D2KO and log_2_FC = 2.6 in D10KO, FDR = 1.5E-259). We validated these findings by qRT-PCR using samples from independent motor neuron cultures (Figure 4D). We also noted a reduction of transferrin (*TF*) expression in KO motor neurons (log_2_FC = −2.8 FDR = 2.5E-125).

To identify cellular pathways that were altered by the loss of CHCHD2 or CHCHD10, we used enrichment analysis (Figure 4E, Table S4). Interestingly, GO term enrichments demonstrated pathways related to neuronal developmental and synapse function, and to ion channel activities, although we had not detected any changes in the ability of the KO motor neurons to fire evoked action potentials. We detected a reduction in several ionotropic glutamate receptor subunits, such as *GRIA1* (AMPA-type receptor subunit), and *GRIK5* (kainate-type receptor subunit), and in NMDA-type glutamate receptor subunits *GRIN2D*, *GRIN3A* and *GRIN3B*, except *GRIN1* that was upregulated. Also several metabotropic glutamate receptor subunits, such as *GRM2*, *GRM3*, *GRM4* and *GRM7*, were upregulated (Figure 4F). In addition, we detected significant changes in the expression of several ion-channel subunits (Figure 4F). K^+^-channel subunits (*KCNK3*, *KCND2*, *KCNJ3*, *KCNIPA* and *KCNT1*), Na^+^ channel subunits (*SCN1A* and *SCN4B*) and Ca^2+^ channel subunits (*CACNG3* and *CACNG8*) were all significantly downregulated. On the contrary, Ca^2+^ channel subunits (*CACNB2*, *CACNA1I* and *CACNA1G*) and Na^+^-channel subunit *SCN1B* were all significantly upregulated.

Finally, we did not detect enrichment of mitochondria-related pathways, indicating that loss of CHCHD2 or CHCHD10 did not result in transcriptional compensation of mitochondrial functions in human motor neurons.

## DISCUSSION

The function of CHCHD10 and CHCHD2 proteins and the mechanisms by which the dominant pathogenic mutations in the corresponding genes cause neurological diseases have been under intense research. Studies have suggested that mice lacking CHCHD10 or CHCHD2 develop normally and show no pathologies (Meng *et al.*, 2017; Burstein *et al.*, 2018; Anderson *et al.*, 2019), supporting gain-of-function mechanism for the dominant patient mutations. CHCHD10 knock-in mice carrying a pathogenic patient mutation, however, died of a severe cardiomyopathy (Anderson *et al.*, 2019; Genin *et al.*, 2019), whereas cardiac phenotypes have not been reported in patients (Penttilä *et al.*, 2017). This suggests that besides mice relevant human cell types should be studied for CHCHD10 and CHCHD2 function and defects. Here we utilized iPSC technology that enabled us to address the importance of CHCHD2 and CHCHD10 in human motor neurons, which are affected in CHCHD10-associated phenotypes. We used CRISPR/Cas9-based gene editing to generate iPSC clones that were lacking either CHCHD2 or CHCHD10, and differentiated those into motor neurons. Knockout motor neurons were compared to neurons differentiated from parental iPSC line that had the identical genome except for the edited gene. Our results showed that iPSC lacking CHCHD2 or CHCHD10 are viable and pluripotent. Furthermore, we demonstrated that CHCHD2 or CHCHD10 are dispensable for human motor neuron differentiation *in vitro*.

We did not observe changes in mitochondrial ultrastructure in D2KO or D10KO iPSC or iPSC-derived motor neurons, nor any signs pointing to compensation of mitochondrial dysfunction, other than increases in CHCHD10 or CHCHD2 amounts. In particular, CHCHD2 absence was compensated for by CHCHD10 increase in both human iPSC and motor neurons. Also a knockout study in HEK293 cells found reciprocal compensatory responses to loss of either protein (Huang *et al.*, 2018). Compensation and functional redundancy, which is also supported by the similar global transcriptome alterations in both knockout motor neurons, may explain why mature motor neurons lacking one of the proteins could be generated. It is, however, possible that double knockout motor neurons would also be functional. We were not successful in generating double knockout iPSC lines in this study and currently are unable to conclude that such iPSC would not be viable. When considering gene therapy options for the dominant CHCHD10/2 mutations, it may be beneficial to know that human motor neurons show reciprocal compensatory responses when the amount of one of the proteins is reduced.

In the transcriptome analysis of CHCHD2/10 knockout motor neurons, we identified highly upregulated expression of *DPP6*, which encodes a single-pass type II membrane protein involved in trafficking of voltage-gated potassium channels (Sun *et al.*, 2011). GWAS associated *DPP6* with susceptibility to sporadic ALS (van Es *et al.*, 2008), and *DPP6* was found as one of the most increased transcripts in iPSC-motor neurons from ALS patients with the *C9ORF72* repeat expansion (Sareen *et al.*, 2013). Another highly upregulated gene in both knockouts was *LYNX1*, which functions as an allosteric modulator of nicotinic acetylcholine receptor, balancing neuronal activity and survival (Miwa *et al.*, 2006). We also saw a reduction of transferrin (*TF*) expression in knockout motor neurons. Transferrin is an extracellular iron transporter, balancing cellular iron uptake with efflux, and it is mainly expressed in liver but also in the nervous system. CHCHD10 knockdown in HEK293 cells was shown to result in accumulation of intramitochondrial iron, leading to a suggestion of CHCHD10 to function as a chaperone in mitochondrial iron import (Burstein *et al.*, 2018). Whether iron levels and transferrin have a role in motor neurons in CHCHD10/2-deficiency remains to be studied.

Although we did not identify alterations in electrophysiological properties of the knockout motor neurons, we found an enrichment of genes in synapse function and synaptic membrane pathways in the transcriptome analysis. We detected upregulation of a subset of metabotropic glutamate receptor subunits, whereas ionotropic subunits were mostly downregulated. Differential expression of glutamate receptor subunit could cause altered synaptic transmission. Even though acetylcholine is the main neurotransmitter at neuromuscular junction (NMJ), there are indications that glutamate might participate in modulating cholinergic transmission at vertebrate NMJ (Colombo & Francolini, 2019). Glutamate activates presynaptic ionotropic receptors leading to increased release of acetylcholine (Fu *et al.*, 1995; Liou *et al.*, 1996). In addition, the expression levels of several K^+^ -, Na^+^- and Ca^2+^-channel transcripts were altered. Altered excitability has been proposed as one of the underlying causes for ALS (Wainger *et al.*, 2014; Naujock *et al.*, 2016). Combining our electrophysiological and transcriptome findings, we propose that although functional human motor neuron differentiation *in vitro* does not require CHCHD2 or CHCHD10, the absence of these proteins does affect the regulation of synaptic function, as detected by transcriptional alterations. Whether this effect is large enough to compromise function for example in a late-onset manner, or in combination with a postsynaptic defect, is not yet known.

Interestingly, a new muscle-specific CHCHD10 knockout mouse showed motor defects, abnormal neuromuscular transmission and NMJ structure, suggesting that CHCHD10 has an important role at the peripheral synapse (Xiao *et al.*, 2019). Similarly, the CHCHD10 knockin mice had severe NMJ degeneration and motor behavior defects (Anderson *et al.*, 2019; Genin *et al.*, 2019). Co-cultures of iPSC-derived motor neurons and myotubes could be used to verify whether human NMJ formation is also affected by CHCHD10/2 defects. It is also possible that both the motor neurons and muscle cells contribute to the dying-back motor neuronopathy, as was reported in the case of FUS-ALS, which revealed endplate maturation defects (Picchiarelli *et al.*, 2019). Future studies using more complex culture systems, which better mimic the physiological environment and interactions of motor neurons, are required to address the consequences that the identified synaptic alterations may have in deficiency of CHCHD2 and CHCHD10.

In summary, we have presented functional differentiation of human iPSC-derived motor neurons in the absence of CHCHD10 or CHCHD2, with reciprocal compensation. Similar transcriptome profiles of the two knockouts support functional redundancy for the two proteins in motor neurons, and the gene expression changes related to synapse functions suggest that loss of CHCHD10 or CHCHD2 is not fully devoid of consequences.

## MATERIAL AND METHODS

### Ethical approval

The generation of the human induced pluripotent stem cell lines used in this study was approved by the Coordinating Ethics Committee of the Helsinki and Uusimaa Hospital District (Nro 423/13/03/00/08) with informed consent of the donor.

### Induced pluripotent stem cells (iPSC)

The reprogramming of human neonatal foreskin fibroblasts from healthy control (HEL46.11) into pluripotent stem cells by Sendai virus technology has been done at Biomedicum Stem Cell Center (University of Helsinki, Finland), as previously described (Trokovic *et al.*, 2015), and the pluripotency confirmed in previous studies (Saarimäki-Vire *et al.*, 2017; Weltner *et al.*, 2018). Using the HEL46.11 iPSC line as the parental control, we generated one D2KO clone and four D10KO clones (F3, F2, G4 and B9), of which we used F3, F2 and G4 in most of the studies. IPSC were maintained on Matrigel-coated (Corning) plates with E8-medium (Gibco) supplemented with E8-supplement (Gibco). IPSC were passaged with 0.5 mM EDTA (Invitrogen) in phosphate-buffer saline (PBS) when confluent. Pluripotency of the generated knockout iPSC lines was confirmed with immunostaining with Nanog (1:250 dilution, D73G4, Cell Signaling) and TRA-1-81 (1:100, MA1-024, ThermoFisher) antibodies. In addition, expression levels of *OCT4* and *SOX2* were analyzed by qRT-PCR.

### Genome editing of iPSC

CRISPR-*Sp*Cas9 technology was used to generate D2KO and D10KO iPSC. GuideRNA (gRNAs) sequences were designed using web-based tools from http://crispr.mit.edu (Hsu *et al.*, 2013) and https://benchling.com (Benchling [Biology Software],2018). The gRNAs were designed to target 5’UTR and the first exon. Transcriptional units for gRNA expression were prepared by PCR amplification and further concatenated using Golden Gate 249
assembly as described in (Balboa *et al.*, 2015), detailed protocol is provided by Addgene (http://www.addgene.org/78311/). Different gRNA pairs were first tested by co-transfection with wild type spCas9 expressing the plasmid CAG-Cas9-T2A-EGFP-ires-puro (Addgene plasmid #73811) into HEK293 cells. The genomic region flanking gRNA target sites was amplified with a pair of gene specific primers and editing efficiency was detected by agarose gel electrophoresis.

Sequences (5’-TGGAAGAGCAGGACGTCACG-3’) and (5’-GTGCGGCTTCGGCTTCCACG-3’) in the 5’UTR and first exon, respectively, were chosen for the CRISPR-SpCas9 targeting sites in *CHCHD2*. Similarly, gRNAs used to create *CHCHD10* knockout were (5’-TCTGTTAGGACCACCGCAGA-3’) and (5’-GCCGCGCTGCGGCTTCCCCG-3’). Genome editing was done essentially as described in (Saarimäki-Vire *et al.*, 2017). Briefly, two million HEL46.11 cells were electroporated with 3 μg of CAG-Cas9-T2A-EGFP-ires-puro endotoxin-free plasmid and 500 ng of each gRNA (Neon Transfection System, 1100 V, 20 ms, two pulses, Thermo Fisher Scientific). Electroporated cells were immediately plated onto prewarmed Matrigel-coated plates containing E8-medium with 10 μM ROCK inhibitor (ROCKi) (Y-27632 2HCl, Selleckchem). Alive, singlet cells positive for GFP fluorescence were sorted on 96-well Matrigel-coated plates containing E8 medium supplemented with 5 μM ROCKi and CloneR™ (StemCell Technologies) using BDInflux Flow Cytometer at the Biomedicum Flow Cytometry Unit (University of Helsinki, Finland). Expandable iPSC colonies were screened by PCR followed by sequencing and subsequently by western blotting.

### Motor neuron differentiation

To differentiate iPSCs into motor neurons, we used the protocol by (Maury *et al.*, 2015; Guo *et al.*, 2017) with slight modifications. Briefly, to obtain neuroepithelial stem cells, iPSC were dissociated with Accutase (Gibco) into small clusters and resuspended in E8 medium containing 5 μM Y-27632 (Selleckchem). The following day, medium was changed to Neuronal basal medium (DMEM F12, Neurobasal vol:vol, with N2 (Life Technologies), B27 (Life technologies) and L-ascorbic acid 0.1 mM (Santa Cruz) supplemented with 40 μM SB431542 (Merck), 0.2 μM LDN-193189 (Merck/Sigma), 3 μM CHIR99021 (Selleckchem), and 5 μM Y-27632 (Selleckchem). From day 3 on, 0.1 μM retinoic acid (Fisher) and 0.5 μM SAG (Calbiochem) was added to neurobasal medium. From day 7 on, BDNF (10 ng/ml, Peprotech) and GDNF (10 ng/ml, Peprotech) were added. 20 μM DAPT (Calbiochem) was added on day 9. Motor neuron progenitor spheroids were dissociated into single cells for plating on day 11 by using Accumax (Invitrogen). Motor neuron progenitors were subsequently plated on poly-D-lysine 50 μg/ml (Merck Millipore) and laminin 10 μg/ml (Sigma-Aldrich) coated plates at 5 × 10^4^ cells per cm^2^. At day 14 retinoic acid and SAG were removed from media. From day 16 on, the cells were switched to motor neuron maturation medium supplemented with BDNF, GDNF, and CNTF (each 10 ng/ml, Peprotech). Media were changed every other day by replacing half of the medium. Motor neurons were ISL1-positive around day 30 of differentiation, after which they were considered mature. Motor neurons were used in experiments at week 5, except at week 7 in patch clamp.

### Immunoblotting

For protein analysis iPSC and iPSC-derived motor neurons were manually collected by scraping on ice and samples were maintained at −80°C until further processing. For western blot, total protein from motor neuronal and iPSC cultures were harvested with RIPA buffer (Thermo Fisher) supplemented with Halt™ protease inhibitor (Thermo Scientific). Protein concentration was quantified with BCA assay (Pierce) according to manufacturer’s instructions. 10 μg of protein/sample was run on TGX 10% and 4-20% acrylamide gels (Bio-Rad) and transferred to 0.2 μm PVDF-membrane (Bio-Rad) with the TransBlot Turbo transfer system (Bio-Rad). Membrane was blocked with 5% milk/TBS-T for 1h RT. The primary and secondary antibodies are listed in supplementary materials. Signal was detected with ECL reagent (Thermo Scientific) and membranes were imaged with Chemidoc XRS+ (Bio-Rad). Image Lab Software (BioRad) was used to quantify the band intensities. For quantification, band intensities were normalized to GAPDH or HSP60.

Assembly of mitochondrial respiratory chain complexes were investigated using Blue-Native-PAGE in mitochondria extracted from iPSC as described by (Konovalova, 2019), using primary and secondary antibodies listed in supplementary materials.

### Quantitative PCR (qPCR) based determination of mitochondrial DNA copy number

DNA was extracted with Macherey-Nagel™ NucleoSpin™. 25 ng of DNA was used for the qPCR reaction. Levels of mitochondrial tRNA-LEU (UUR) (*MT-TL1*) and nuclear *B2M* were analyzed by qPCR amplification in CFX Real-time system C1000Touch (Bio-Rad) with SYBR-green Flash (Thermo Fisher) by transcript specific primers. The ratio between *MT-TL1* and *B2M* transcripts were used to determine the copy number of mtDNA. The ratio between mitochondrial cytochrome B (*MT-CYTB*) or nuclear *ACTB* transcripts were used to verify results.

### Immunocytochemistry

IPSC and iPSC-derived motor neuronal cultures on cover glasses were fixed with 4% paraformaldehyde (PFA) for 15 min at RT and then permeabilized with PBS containing 0.2% Triton X-100 (Fisher) for 10 min. Cells were blocked with 5% protease-free BSA (Jackson Immuno Research) in 0.1% Tween20 in PBS (PBS-T) for 2 h in RT. Cells were incubated overnight at +4°C in blocking buffer (5% bovine serum) containing different primary antibodies, listed in supplementary materials. After washing with PBS-T, cells were incubated with secondary antibody for 1h at RT. Cover glasses were applied on microscope slides with Vectashield DAPI (#H-1200) and imaged with Axio Observer Z1 (Zeiss) inverted fluorescence microscope.

IPSCs with CHCHD2/10 (Alexa 488) staining were imaged with LSM880 IndiMo Axio Observed laser scanning confocal microscope with Plan-Apochromat 63x/1.4 oil DIC M27 immersion lense. CHCHD2/10 channel was captured with 488 laser using the same settings and exposure time for all samples (laser 3.06%). Images were acquired at 16-bit depth and 1912-x1912 frame size. The results were visualized with Zen Blue software.

### Quantitative Reverse Transcription PCR (qRT-PCR)

RNA from differentiated motor neuronal cultures was isolated using NucleoSpin RNA extraction kit (Macherey-Nagel). RNA was reverse transcribed with Maxima first strand cDNA synthesis kit (Thermo Fisher). cDNA levels were analyzed by qRT-PCR amplification in CFX Real-time system C1000T (Bio-Rad) with SYBR green Flash (Thermo Fisher) by transcript specific primers. Technical duplicates were used for all studies. 2 ng of cDNA was used for total mRNA quantification. Relative quantification was done using the ΔΔCt method with normalization to *GAPDH*. A list of primers can be found at supplementary methods.

### Transmission electron microscopy (TEM)

Motor neuronal cultures on glass slides/cover glasses were fixed with 2% glutaraldehyde in 0.1 M NaCac buffer for 30 min at RT, and then washed with NaCac buffer. Cells were kept in NaCac buffer at +4°C until further use. iPSC on glass slides were fixed with 2% PFA + 2% Glutaraldehyde (Sigma) for 30 min at 4°C, then washed once with PBS and fixed with 2% Glutaraldehyde O/N at 4°C. Fixed neurons and iPSC were prepared according to standard protocols for transmission electron microscopy. Ultrathin sections were observed with Jeol 1400 transmission electron microscope at 80 000 V. ImageJ (https://imagej.nih.gov/ij/) package Fiji (Schindelin *et al.*, 2012) was used to assess mitochondria length.

### RNA sequencing

RNA was isolated from four independent motor neuron cultures per knockout cell line (D2KO and D10KO G4) and from eight control cultures at day 37 of differentiation. RNA was extracted with Nucleospin RNA kit (Macherey-Nagel) according to manufacturer’s instructions including rDNAse treatment. RNA concentration was measured with Qubit. RNA quality determination and mRNA sequencing were done at Biomedicum Functional Genomics Unit with single-end sequencing with poly(A) binding beads and NEBNext Ultra Directional RNA Library Prep and sequenced with Illumina NextSeq High Output 1 × 75bp.

Raw data was demultiplexed with bcl2fastq2 (v2.20.0.422; Illumina). Adapter sequences (AGATCGGAAGAGCACACGTCTGAACTCCAGTCA), low quality (Phred score<25) and ambiquous bases (N) were removed with cutadapt (v.2.2) (Martin, 2011), retaining reads longer than 25bp. The trimmed reads were mapped to the human reference genome (GRCh38) with STAR (v. 2.5.3) (Dobin *et al.*, 2013). Gene expression was analyzed with R (v.3.6.0) (R Core Team, 2019). The read counts for genes (mapping to exons) were extracted with Rsubreads (v.1.34.6) (Liao *et al.*, 2019) excluding duplicates, multimapping reads, chimeric fragments, and reads with mapping quality below 10.

Differential gene expression was analyzed with DESeq2 (v.1.24) (Love *et al.*, 2014), comparing D2KO and D10KO to parental cells. The differentially expressed genes were analyzed for pathway enrichment using the GO (Ashburner *et al.*, 2000; The Gene Ontology Consortium, 2018), using ClusterProfiler (v3.12.0) (Yu *et al.*, 2012). For enrichment analysis, the set of background genes (universe) was limited to genes expressed in our samples. Genes with low read depth (<50 reads in total) were excluded from the differential expression and enrichment analysis.

RNA-seq data from the motor neurons in this study were contrasted to publicly available datasets of other human iPSC derived motor neurons; GSE108094 (Rizzo *et al.*, 2019) and GSE121069 (Nijssen *et al.*, 2018) as well as laser captured motor neurons; GSE76220 (Batra *et al.*, 2016), GSE76514 (Nichterwitz *et al.*, 2016) and GSE93939 (Allodi *et al.*, 2019). The same analysis procedures (trimming, mapping and read counting) were applied to the reference samples as for our dataset. Gene expression results were visualized with ggplot2 (v3.2.1) (Wickham, 2016), limma (v3.40.6) (Ritchie *et al.*, 2015) and eulerr (v5.1.0) (Larsson, 2019).

### Electrophysiology

Whole-cell patch-clamp recordings were done on living cells after 42 days in culture. Coverslips containing cultured neurons were placed to recording chamber on the visually guided patch-clamp setup. Temperature at the recording chamber was kept at 30°C, and cells were continuously perfused with ACSF of the following composition (mM): 124 mM NaCl, 3 mM KCl, 1.25 mM NaH_2_PO_4_, 1 mM MgSO_4_, 26 mM NaHCO_3_, 2 mM CaCl_2_, 15 mM glucose, and 95 % O_2_/5 % CO_2_. Recordings were done with glass microelectrodes (4-6 MOhm), using Multiclamp 700A amplifier (Axon Instruments, USA), and digitized at 20 kHz. Microelectrodes were filled with the intracellular solution containing (mM): 105 K gluconate, 15 KCl, 5.3 CaCl_2_, 5 NaCl, 10 HEPES, 10 EGTA, 4 MgATP, 0.5 Na_2_GTP, pH 7.2. Data were collected with Clampex 10.2 (Molecular Devices, USA).

Series resistance, input resistance and capacitance were recorded in voltage clamp mode at −70 mV by injection of 5 mV voltage steps. If series resistance exceeded 20 MΩ, or if the change in series resistance during the experiment exceeded 30 %, the recording was discarded.

Voltage-gated sodium/calcium and potassium currents were recorded in voltage clamp mode. Baseline voltage was set at −70 mV, and voltage steps with the increasing increment of 10 mV were injected for 100 ms. Leak subtraction protocol was applied, and the current amplitude was analyzed after correction for leak current. Resting membrane potential and spontaneous activity were recorded in current clamp mode without any background current injected. For recording of the evoked action potentials, baseline membrane potential was held at −60 mV, and depolarizing current steps with the increasing increment of 10 pA were injected for one second.

Data were analyzed in Clampfit 10.2 (Molecular Devices, USA). Properties of individual evoked action potentials were analyzed by a custom MATLAB script (MATLAB R2018b, MathWorks, USA; Zubarev, Nagaeva, unpublished). Meta-analysis was done in GraphPad Prism 8.

### Statistical analysis

GraphPad Prism was used for all statistical analyses except RNA-seq. Unless otherwise indicated, mean ± SD values are reported in the graphs. One-way ANOVA was used for the experiments with post-hoc Tukey’s test to determine statistical differences between groups. *P < 0.05 were considered significant.

## Data availability

Differential expression analysis and raw data for the RNA-Seq are deposited at NCBI Gene Expression Omnibus under the accessions GSE133764 (CHCHD2 KO) and GSE133763 (CHCHD10 KO).

## Acknowledgments

We thank Riitta Lehtinen for technical assistance. We acknowledge Biomedicum Stem Cell Centre, Biomedicum Functional Genomics Unit, Biomedicum Imaging Unit, and Electron Microscopy Unit of the Institute of Biotechnology, University of Helsinki. HiLIFE Flow cytometry unit is acknowledged for assistance with cell sorting. This work was supported by the Academy of Finland (MetaStem Centre of Excellence to H.T., Clinical Researcher Funding to E.Y.), Sigrid Juselius Foundation, University of Helsinki, Emil Aaltonen Foundation, and Helsinki University Hospital.

## Author Contributions

Conceptualization, S.H., R.W., E.Y., H.T.; Methodology and Investigation S.H., T.S.R., S.M.M., R.W., S.K., M.S.T., J.P., H.I.; Formal analysis, S.H. and J.K.; Writing – Original Draft Preparation S.H. and H.T.; Writing – Review & Editing, all authors; Project Administration H.T.; Funding Acquisition E.Y. and H.T.; Supervision, T.T., T.O., E.Y. and H.T.

