## Supplemental text and figures for "Chchd10 or Chchd2 are not Required for Human Motor Neuron Differentiation *In Vitro* but Modify Synaptic Transcriptomes"

**Supplementary materials**

**Table S1. Antibody list.**

| Antibodies | | |
|---|---|---|
| ChAT | Millipore | Cat#AB144P, RRID:AB_2079751 |
| CHCHD10 | Sigma-Aldrich | Cat#HPA003440, RRID:AB_1078348 |
| CHCHD2 | Sigma-Aldrich | Cat#HPA027407, RRID:AB_10959659 |
| Complex I (NDUFA9) | Abcam | Cat# ab14713, RRID:AB_301431 |
| Complex II (SDHA) | Abcam | Cat# ab14715, RRID:AB_301433 |
| Complex III (UQCRC2) | Abcam | Cat# ab14745, RRID:AB_2213640 |
| Complex IV (COX1) | Abcam | Cat# ab14705, RRID:AB_2084810 |
| Complex V (ATP5A) | Abcam | Cat# ab14748, RRID:AB_301447 |
| GAPDH | Cell Signaling Technology | Cat# 5014, RRID:AB_10693448 |
| HB9 | DSHB | Cat# 81.5C10, RRID:AB_2145209 |
| HSP60 | Santa Cruz Biotechnology | Cat# sc-1052, RRID:AB_631683 |
| ISL1 | DSHB | Cat# 39.4D5, RRID:AB_2314683 |
| MAP2 | Millipore | Cat# AB5543, RRID:AB_571049 |
| Nanog | Cell Signaling Technology | Cat# 4903, RRID:AB_10559205 |
| NEFH (SMI-32) | Biolegend | Cat# 801701, RRID:AB_2564642 |
| NEFM | Proteintech | Cat# 20664-1-AP, RRID:AB_10700009 |
| TRA-1-81 | Thermo Fisher Scientific | Cat# MA1-024, RRID:AB_2536706 |
| TUJ1 | BioLegend | Cat# 801201, RRID:AB_2313773 |

**Table S2. Primers for qPCR.**

| Name | Forward primer sequence 5’-3’ | Reverse primer sequence 5’-3’ |
|---|---|---|
| *B2M* | TGCTGTCTCCATGTTTGATGTATCT | TCTCTGCTCCCCACCTCTAAGT |
| *ACTB* | CCTGGCACCCAGCACAAT | GGGCCGGACTCGTCATAC |
| *CHAT* | GGAGGCGTGGAGCTCAGCGACACC | CGGGGAGCTCGCTGACGGAGTCTG |
| *CHCHD10* | GTGACCTGTCCCTGTGTGAG | TCTGTGGGGTGAGAAACCTC |
| *CHCHD2* | AGGGTTTCAATGAGGTGCTG | GAAGCCAGGAATGACAGGAG |
| *MT-CYB* | GCCTGCCTGATCCTCCAAAT | AAGGTAGCGGATGATTCAGCC |
| *GAPDH* | CGCTCTCTGCTCCTCCTGTT | CCATGGTGTCTGAGCGATGT |
| *ISL1* | GTTACCAGCCACCTTGGAAA | GGACTGGCTACCATGCTGTT |
| *MAP2* | CCAATGGATTCCCATACAGG | CTGCTACAGCCTCAGCAGTG |
| *NEFM* | TGCAGTCCAAGAGCATCGAGC | AGTCTCTTCACCCTCCAGGAGTT |
| *MT-TL1* | CACCCAAGAACAGGGTTTGT | TGGCCATGGGTATGTTGTTA |
| *TUBB3* | GCCAAGTTCTGGGAAGTCAT | CCACTCTGACCAAAGATGAA |

**Table S3. Differential expression analysis.** Comparisons between knockout (KO) and wild type (WT) for CHCHD2 and CHCHD10, and comparison between the two knockouts. Results are grouped and colored the same way as in Venn diagram (Figure 4C).

**Table S4. GO term enrichment analysis.** Results of differentially expressed genes between knockout (KO) and wild type (WT) for CHCHD2 and CHCHD10. Genes in the universe are limited to genes expressed in the samples (total read counts > 50).

### Supplementary figures

**Figure S1. Successful generation of knockout iPSC lines.** (A) Expression of *CHCHD2* and *CHCHD10* in knockout iPSCs by qRT-PCR, normalized to *GAPDH*, individual D10KO clones shown. F3 and F2 show residual ~10% expression of CHCHD10 (n=3 per clone). Data are shown as mean ± SD. **(**B) Uncropped images of the immunoblot of CHCHD10 of whole iPSC lysates. 14kDa CHCHD10 as well as higher molecular weight bands are absent in all D10KO clones. Mitochondrial HSP60 shown as a loading control. (C) Uncropped images of the immunoblot of CHCHD2 of whole iPSC lysates. Immunoblotting with polyclonal CHCHD2 antibody shows co-immunoreactivity with highly up-regulated CHCHD10, indicated by asterisks. Mitochondrial HSP60 shown as a loading control.

**Figure S2. Increased CHCHD10 in D2KO motor neurons. (**A) Uncropped images of the immunoblot with CHCHD10 show increased 14kDa CHCHD10 and higher molecular weight bands in D2KO neurons that are absent in all D10KO clones. (B) Uncropped images of the immunoblot with CHCHD2 antibody. GAPDH is shown as a loading control.

**Figure S3. Characterization of neuronal and motor neuronal identity in CHCHD2/10 knockout neurons compared to parental control. (**A) Immunocytochemical analysis of TUJ1 in neuronal culture. Scale bar 50 µm. (B) Immunocytochemical analysis of motor neuronal non-phosphorylated neurofilament heavy, NEFH (SMI-32) (red) and acetylcholine transferase (ChAT) (green) in neuronal culture confirming succession of motor neuron differentiation of D2KO and D10KO iPSCs. Dapi indicates nuclear staining. Scale bar 50 µm.

Supplemental figure 1.

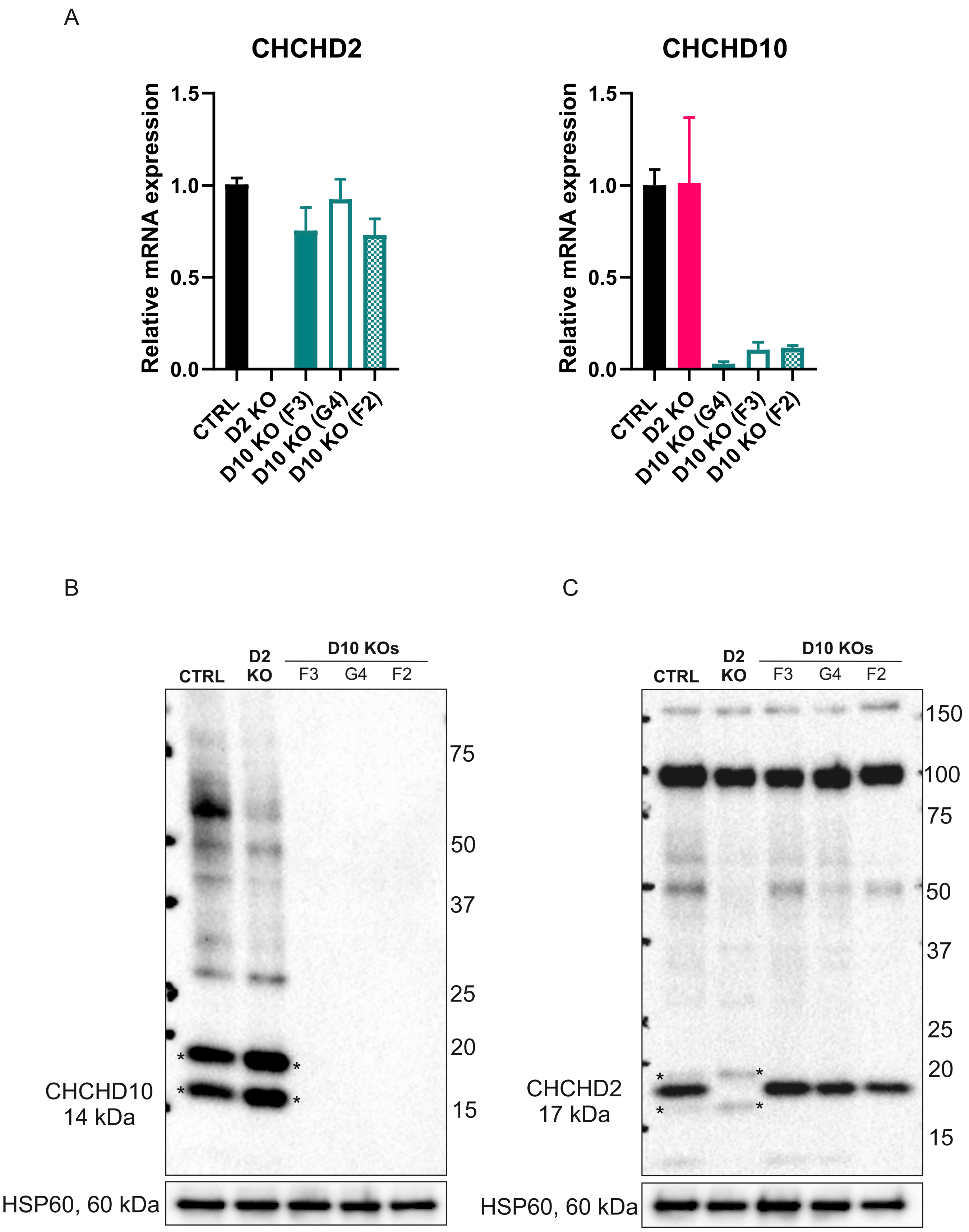

Supplemental figure 2.

A

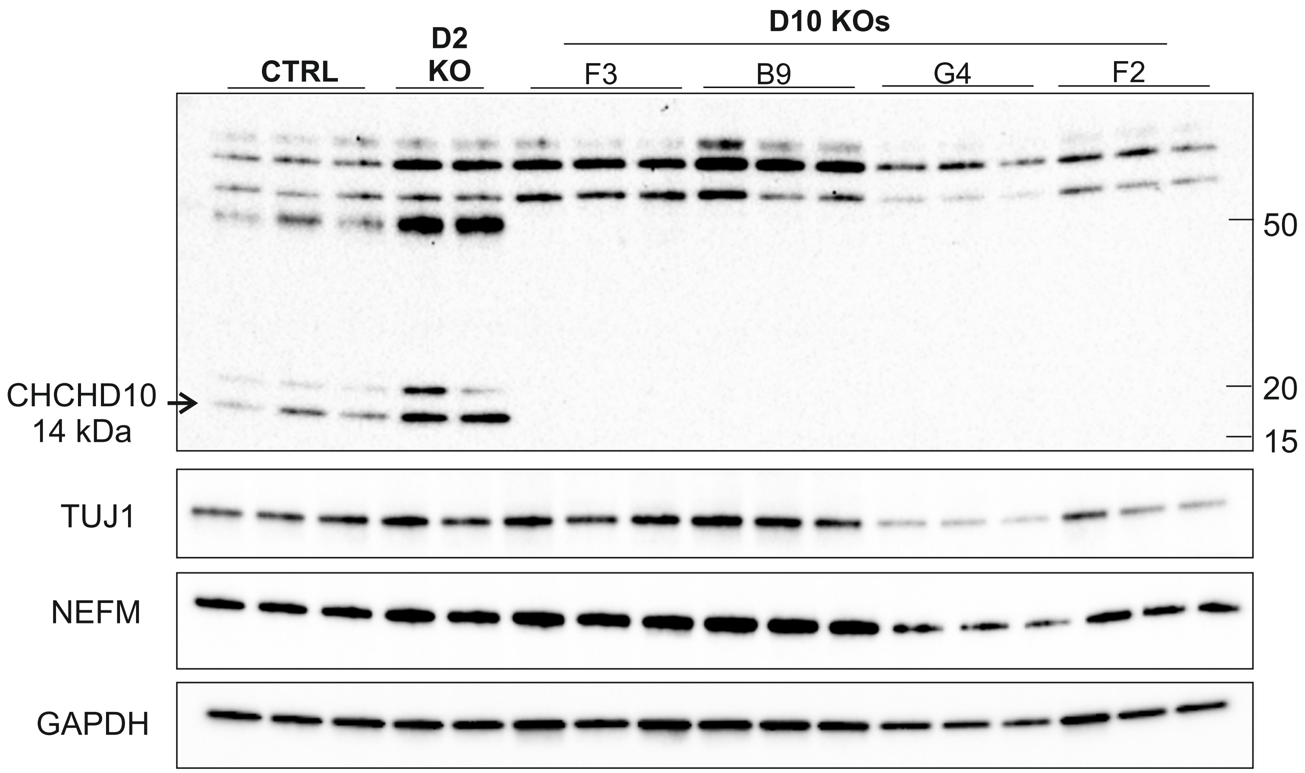


B

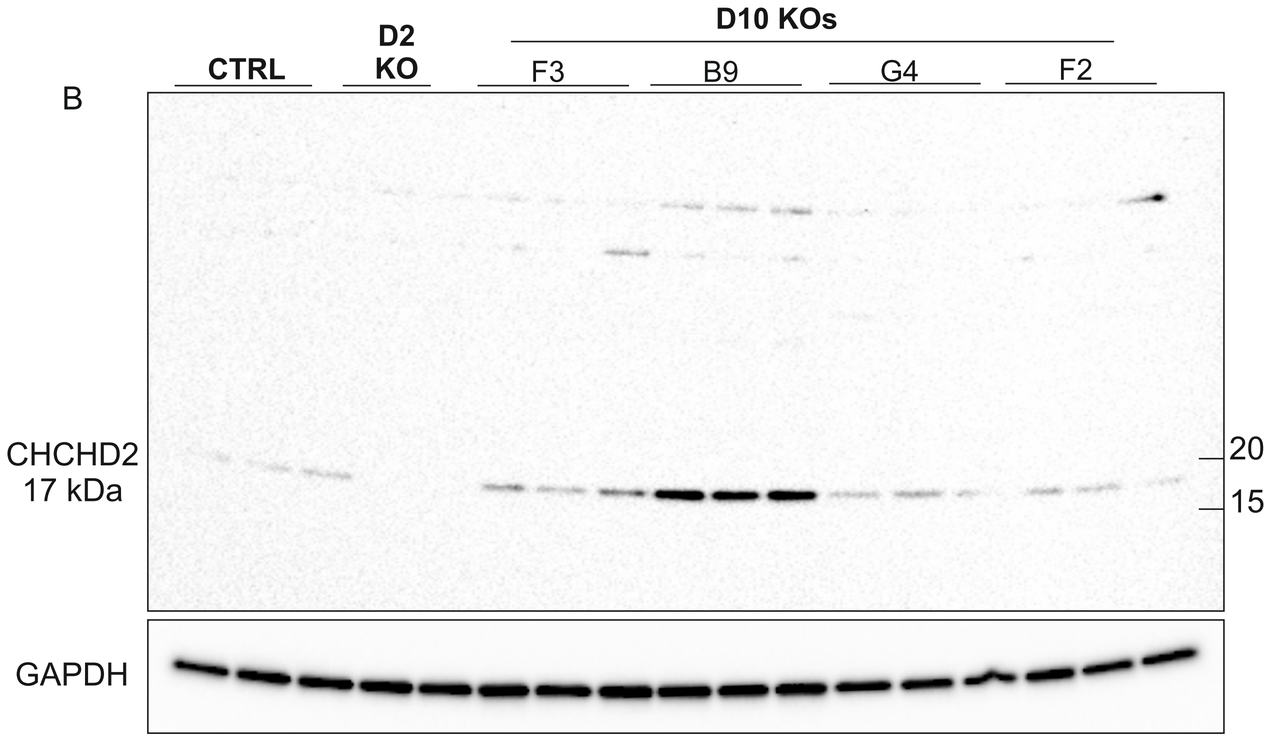

Supplemental figure 3.

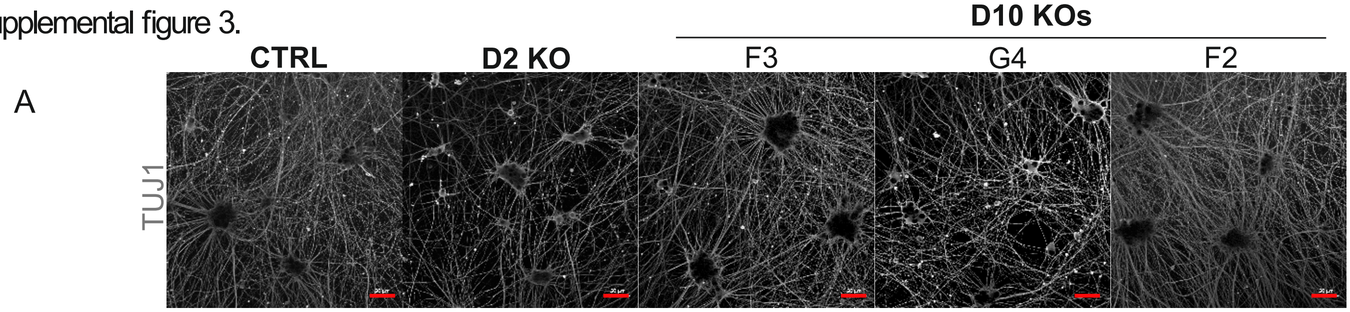


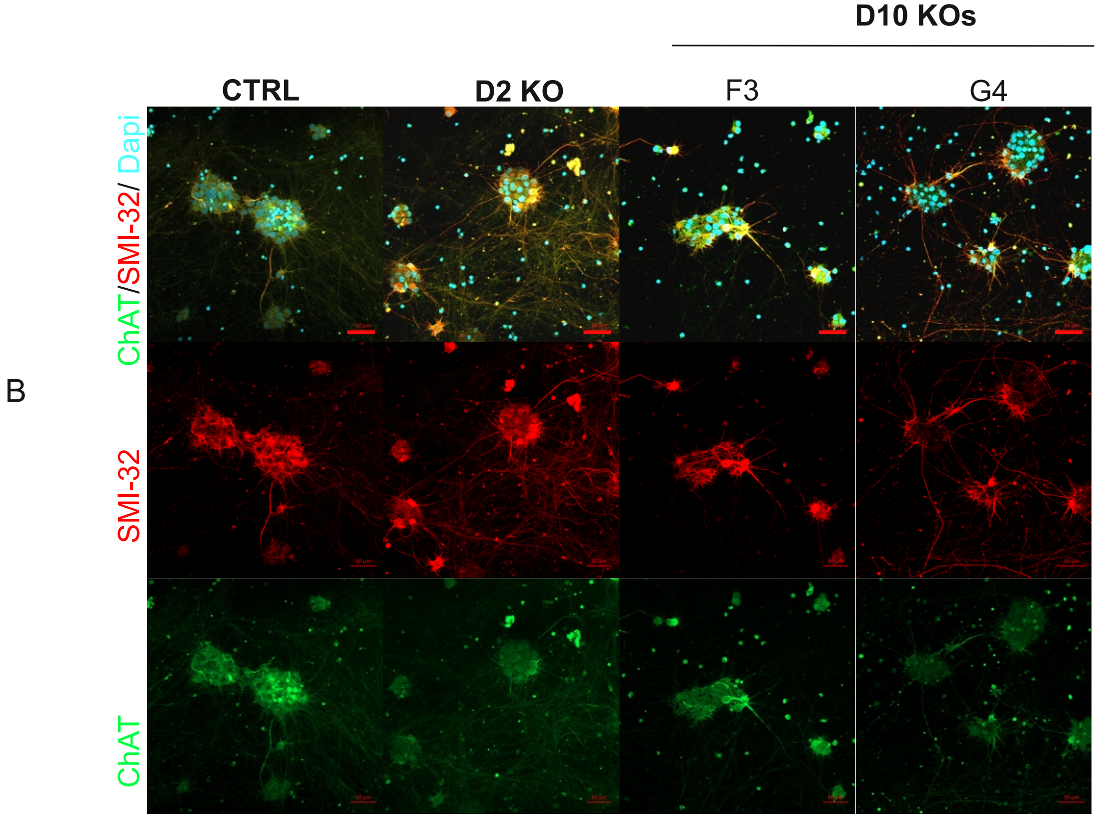
